# Analysis of the oxidative stress regulon identifies *soxS* as a genetic target for resistance reversal in multi-drug resistant *Klebsiella pneumoniae*

**DOI:** 10.1101/2020.08.21.262022

**Authors:** João Anes, Katherine Dever, Athmanya Eshwar, Scott Nguyen, Yu Cao, Sathesh K Sivasankaran, Sandra Sakalauskaitė, Angelika Lehner, Stéphanie Devineau, Rimantas Daugelavičius, Séamus Fanning, Shabarinath Srikumar

## Abstract

In bacteria, the defense system deployd to counter oxidative stress is orchestrated by three transcriptional factors – SoxS, SoxR, and OxyR. Although the regulon that these factors control is known in many bacteria, similar data is not available for *Klebsiella pneumoniae*. To address this data gap, oxidative stress was artificially induced in *K. pneumoniae* MGH 78578 using paraquat and the corresponding oxidative stress regulon recorded using RNA-seq. The *soxS* gene was significantly induced during oxidative stress and a knock-out mutant was constructed, to explore its functionality. The wild-type and mutant were grown in the presence of paraquat and subjected to RNA-seq to elucidate the *soxS* regulon in *K. pneumoniae* MGH78578. Genes that are commonly regulated both in the oxidative stress regulon and *soxS* regulon were identified and denoted as the ‘oxidative SoxS regulon’ – these included a stringent group of genes specifically regulated by SoxS. Efflux pump encoding genes such as *acrAB-tolC, acrE*, and global regulators such as *marRAB* were identified as part of this regulon. Consequently, the isogenic *soxS* mutant was found to exhibit a reduction in the minimum bactericidal concentration against tetracycline compared to that of the wild type. Impaired efflux activity, allowing tetracycline to be accumulated in the cytoplasm to bactericidal levels, was further evaluated using a tetraphenylphosphonium (TPP^+^) accumulation assay. The *soxS* mutant was also susceptible to tetracycline *in vivo*, in a zebrafish embryo model. We conclude that the *soxS* gene could be considered as a genetic target against which an inhibitor could be developed in the future and used in combinatorial therapy with tetracycline to combat infections associated with multi-drug resistant *K. pneumoniae*.

## Introduction

Oxygen started accumulating in the biosphere about 2-3 billion years ago. Many organisms harvest energy by oxidizing organic compounds with oxygen acting as the terminal electron acceptor. This molecule, therefore, has become essential for life, at least for aerobic organisms. As a natural consequence of aerobic metabolism, the production of toxic reactive oxygen species (ROS) namely, hydrogen peroxide (H_2_O_2_), superoxide radical (O_2_^•-^), and the generation of hydroxyl radical (HO^•^), becomes inevitable in an oxygen-rich environment. Different ROS will not only oxidize macromolecules (such as DNA, proteins, lipids) but also extract iron from proteins containing an iron-sulfur cluster creating a highly reactive HO^•^ rich intracellular environment ^1,2^, detrimental for bacteria. Therefore, to survive the effects of ROS, bacteria deploy a variety of adaptive responses. These are well characterized in bacteria like *Escherichia coli* ^3^, but as yet, not in *Klebsiella pneumoniae*

In bacteria, the primary antioxidant defence systems employ superoxide dismutase (SOD) and catalase (CAT) enzymes ^1,4^. However, these systems may prove inadequate to protect bacteria under circumstances of extreme and prolonged oxidative stress. During these stress conditions, bacteria can activate the OxyR and SoxRS systems in response to hydrogen peroxide ^5^ and redox-active compounds ^6^, respectively. Both the OxyR and SoxRS work by transcriptionally activating genes whose protein products either protect or repair the damage caused by intracellular ROS accumulation. In the SoxRS system, the activation of a target gene occurs *via* a two-step process wherein SoxR acts as a sensory protein recognizing elevated levels of ROS. Under normal conditions (non-stressed), the binuclear iron-sulfur clusters [2Fe-2S] in the SoxR protein remain reduced. In the presence of enhanced levels of superoxides, the [2Fe-2S] clusters are oxidized ^7^. Oxidization of the SoxR protein enhances the open complex formation of SoxR with RNA polymerase, thereby activating the transcription of *soxS* ^8^. The SoxS protein is a transcriptional activator belonging to the XylS/AraC family ^7^. SoxR dependent induction of SoxS, in turn, activates the transcription of many other genes (denoted collectively as the SoxRS regulon) whose primary functions involve anti-oxidative action, detoxification, efflux of redox-active compounds, changes in membrane permeability and protecting DNA thereby rescuing bacteria from the deleterious effects of increased intracellular levels of ROS ^9-11^. In *E. coli* genes that were regulated by SoxS were identified ^12-17^. Overall, the biological role of the SoxRS regulon can be summarized as (1) prevention of oxidative damage (2) recycling of damaged macromolecules and (3) regeneration of nicotinamide adenine dinucleotide phosphate.

Though the transcriptional organization of *soxS* is well characterized in some pathogens, these data are lacking for *K. pneumoniae*. These bacteria are a member of the ESKAPE group, one of six pathogens responsible for most drug-resistant nosocomial infections ^18^. Since oxidative stress is known to mediate antibiotic resistance in pathogens we were interested in identifying how *K. pneumoniae* responds to oxidative stress and what its impact might be on antimicrobial resistance. In this study, RNA-seq was used to describe the transcriptional architecture of *K. pneumoniae* MGH78578 during exposure to a ROS inducing agent, paraquat, revealing that the regulon was controlled by the *soxRS* two-component system. In addition, RNA-seq analysis of the isogenic mutant *K. pneumoniae* MGH 78578Δ*soxS* was carried out and used to describe the ‘oxidative *soxS* regulon’ - a stringent set of genes regulated *via soxS. K. pneumoniae* MGH 78578Δ*soxS* was highly susceptible to tetracycline. Susceptibility of the mutant to tetracycline coupled with increased accumulation of tetraphenylphosphonium (TPP^+^) in the bacterial cytoplasm was supported at least in part by the down-regulation of *acrAB-tolC* and the global regulator *marRAB* in *K. pneumoniae* MGH 78578Δ*soxS*. Since the mutant was highly avirulent in a zebrafish model, we predict that *soxS* can be used as a genetic target to inhibit infections associated with MDR *K. pneumoniae*.

## Results and discussion

### SoxS is the major transcriptional regulator when *K. pneumoniae* MGH78578 is exposed to redox compound-based oxidative stress

Experimentally, oxidative stress can be induced in bacteria by exposing cultures to either redox compounds like PQ (paraquat) or H_2_O_2_. PQ is 1,1-dimethyl-4,4-bipyridinium and is a widely used nonselective herbicide, found to induce oxidative stress by enhancing ROS levels - superoxide anion radical (^•^O_2_ ^-^) in a dose-dependent manner, as exemplified in *Vibrio cholera, E. coli* and others ^14,19^. Firstly, we started by assessing the inhibitory concentration of PQ in *K. pneumoniae* MGH78578 using broth microdilution and determined the MIC to be 15.62 µM. Thereafter, the following transcriptomic experiments were carried out at sub-minimal inhibitory concentrations (half the MIC). Here, we used RNA-seq to investigate the genome-wide transcriptional architecture of multi-drug resistant *K. pneumoniae* MGH78578 following exposure to a sub-inhibitory concentration of PQ. *K. pneumoniae* MGH78578 was exposed to 7.8 µM PQ for 30 minutes to induce oxidative stress. These PQ induced cultures (denoted as MGH_PQ_ A and B) along with a parallel set of unexposed wild type bacterial cells (denoted as MGH_wt_ A and B) were subjected to RNA isolation and deep-level sequencing (Table S1-Work Sheet [WS]1). Similarly, libraries were also prepared using the isogenic *soxS* mutant (denoted as MGH*ΔsoxS*_PQ_ A and B libraries and are discussed elsewhere) constituting the six RNA-seq libraries generated in this study.

Approximately 57 million uniquely mapped reads were generated across all six libraries accounting for more than 9 million reads/library (Table S1-WS1), data that was sufficient for robust transcriptional analysis ^20^. The expression levels of 5,185 *K. pneumoniae* MGH78578 chromosomal genes and the resident plasmid encoding genes (including plasmids pKPN3, pKPN4, pKPN5, pKPN6, and pKPN7) were calculated using the *Voom* approach (limma package) ^21^. We confirmed the reproducibility of the RNA-seq data by calculating the Spearman coefficients for the biological replicates of all libraries based on the normalized read counts. In all six libraries, the coefficient was found to be ∼ 0.96 to 0.99, confirming the statistical significance between replicates (Fig. S1).

Here, we describe the ‘oxidative stress regulon’ of *K. pneumoniae* MGH78578 by identifying the genes that were differentially regulated in MGH_PQ_ *versus* MGH_wt_ libraries. The oxidative stress regulon comprised 1,366 genes which were differentially regulated (Table S1-WS2 and Fig. 1A). Of these 11.5% (n=158) were highly up-regulated (>4-fold) and 22.5% (n=309) were up-regulated (2 to 4-fold). A total of 49 genes (3.7%) were highly down-regulated (>4-fold) while a further 147 (11.12%) were down-regulated (2 to 4-fold) (Table S1-WS2). Upon analysis, the most induced *K. pneumoniae* MGH78578 gene was found to be *soxS* (145-fold) indicating that the *soxRS* regulon was highly active in PQ exposed *K. pneumoniae* MGH78578. The transcriptomic response of bacteria to oxidative stress is specific to the agent causing oxidative stress - extracellular H_2_O_2_ triggers OxyR regulon while PQ induces the SoxRS regulon, as exemplified in *Escherichia coli* ^3^. Based on these *E. coli* data, we hypothesized that exposure to PQ should induce the SoxS regulon in *K. pneumoniae* MGH78578. Since no data was available in *K. pneumoniae*, we put our hypothesis to the test using RT-qPCR targeting the *soxS* gene. Our RT-qPCR data confirmed that the expression of the *soxS* transcript improved with increasing concentration of PQ (Fig. 2A).

**Fig. 1.**
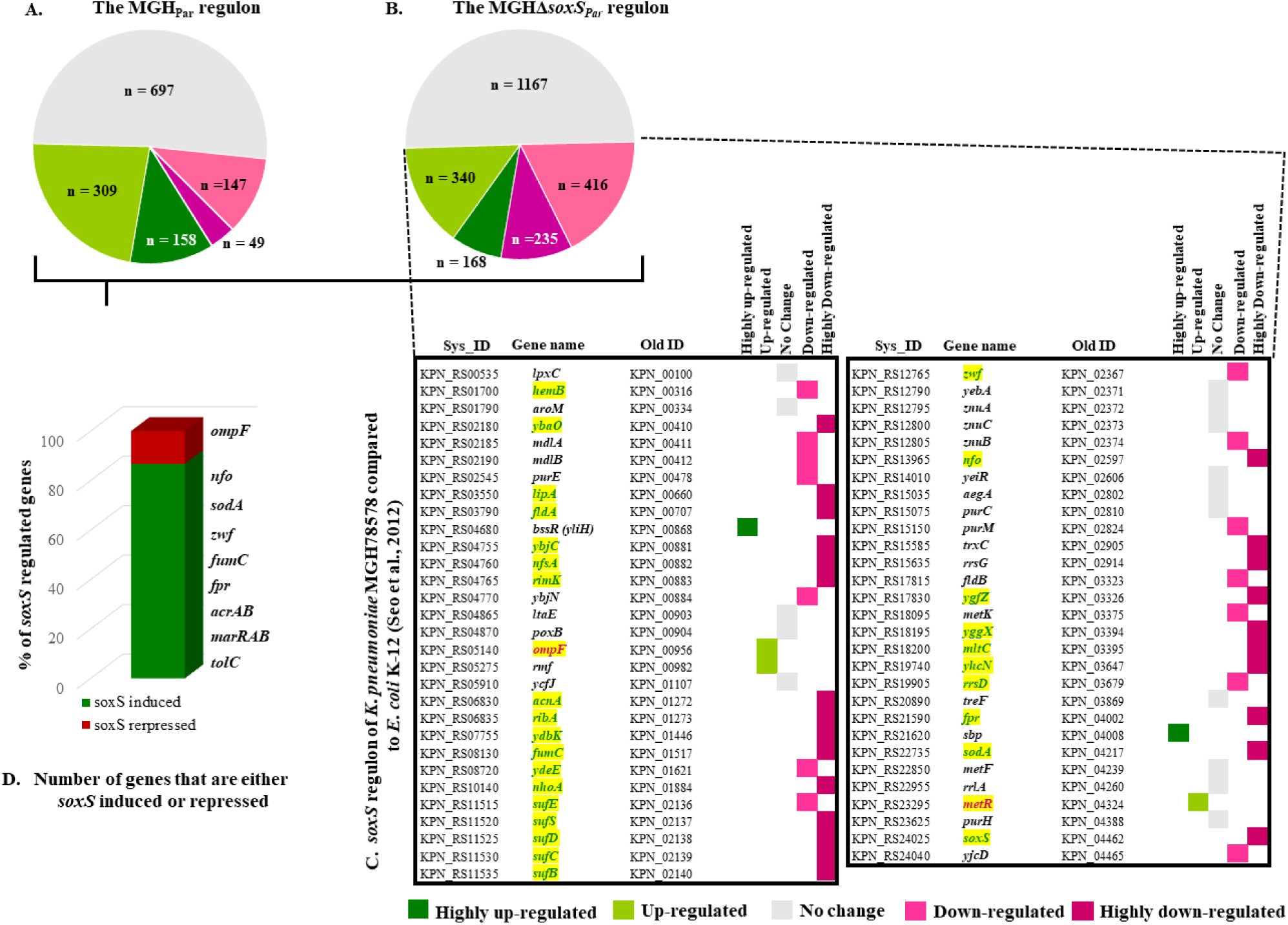
The oxidative, *soxS*, and ‘oxidative *soxS*’ regulon of *K. pneumoniae* MGH78578. **(A)**. The number of statistically significant genes identified in the oxidative regulon of *K. pneumoniae* MGH78578. The number of genes are categorized according to their expression pattern and depicted in a color core based on the color key given below. **(B)**. The number of statistically significant genes identified in the *soxS* regulon and categorized according to their expression pattern. **(C)**. Differentially regulated genes common in the *soxS* regulon of both *K. pneumoniae* MGH78578 and *E. coli* K-12 ^3^. The genes that are in green font and highlighted in yellow are those identified in the ‘oxidative *soxS*’ regulon. **(D)**. The number of genes in the ‘oxidative *soxS*’ regulon of *K. pneumoniae* MGH78578 expressed as a percentage. The most significant *soxS* induced and repressed genes are indicated.

**Fig. 2.**
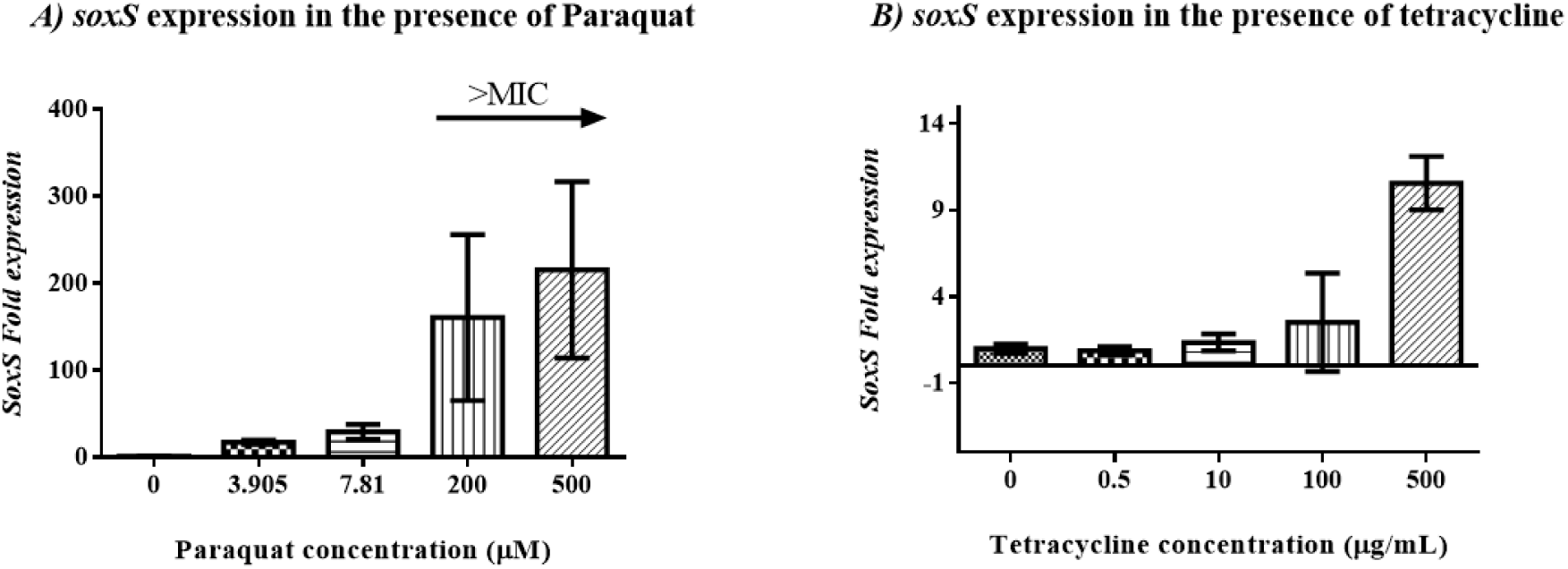
Expression of *soxS* gene under paraquat stress **(A)** and under tetracycline stress **(B)**. In both cases the compounds were added to cells growing at mid exponential phase for 30 minutes.

Exposure to H_2_O_2_, however, generated a different response in other bacteria. Exposure of *V. cholerae* to oxidative stress increased the activity of SOD and CAT enzymes ^19^. However, in *V. cholerae*, the level of CAT did not increase post-exposure to PQ but rather increased during exposure to H_2_O_2_. Our results describing PQ exposed *K. pneumoniae* MGH78578 support this observation – none of the catalases (encoded by genes KPN_RS06170, KPN_RS06615, and KPN_RS09805) were differentially regulated (Table S1-WS2). However, the SOD (encoded by *sodA, sodB*, and *sodC*) was highly up-regulated – *sodA* alone was highly up-regulated (14-fold) while *sodC* was up-regulated (∼ 3-fold) in PQ induced cells. We did not find *sodB* to be differentially regulated within PQ treated *K. pneumoniae* MGH78578. It is tempting to speculate that selective differential regulation of SOD and not CAT in *K. pneumoniae* MGH78578 could be the response to ^•^O_2_ ^-^ induced by PQ.

Since SoxS, a XylS/AraC type transcriptional regulator, was highly induced following exposure to PQ, we were interested in identifying the associated genes that were differentially regulated. For this, we constructed a *K. pneumoniae* MGH78578 Δ*soxS* mutant. We cultured the mutant, exposed the cells to PQ, and again used RNA-seq (MGHΔ*soxS*_PQ_ library) to identify the differentially regulated genes (MGHΔ*soxS*_PQ_ library *versus* MGH_PQ_ library), thus comprising the ‘*soxS* regulon’. The *soxS* regulon was made up of 2,326 differentially regulated genes (Table S1-WS2) (Fig. 1B). Of these 7.2% of the genes (n=168) were highly up-regulated (>4-fold) and 14.6% (n=340) were up-regulated (2-4-fold). A total of 235 genes (10%) were highly down-regulated (>4-fold) while 416 (17.8%) were down-regulated (2-4-fold) (Table S1-WS2) (Fig. 1B). To demonstrate the robustness of these data, we compared our *K. pneumoniae* MGH78578 *soxS* regulon with the *E. coli soxS* regulon published earlier ^3^. Of the 59 *soxS* regulated in *E. coli* K-12 genes, 44 were also found to be similarly regulated by *soxS* in *K. pneumoniae* MGH78578 (Fig. 1C).

To add stringency to our data, we further compared the oxidative stress regulon to the *soxS* regulon to identify those genes that belonged to the ‘oxidative *soxS*’ regulon. The oxidative *soxS* regulon represented a stringent set of *K. pneumoniae* MGH78578 genes that were regulated by *soxS* alone. The genes belonging to this regulon had a characteristic statistically significant expression pattern - up-regulated in the MGH_PQ_ and down-regulated in MGHΔ*soxS*_PQ_ (*i*.*e*. SoxS induced); down-regulated in MGH_PQ_ and up-regulated in MGHΔ*soxS*_PQ_ (*i*.*e*. SoxS repressed). In total, 256 genes belonged to the ‘oxidative *soxS’* regulon. Of these 222 genes were found to be ‘SoxS induced’ while 34 were ‘SoxS repressed’ (Table S1-WS3). Examples include *soxS, acrAB, tolC* among others, all of which were *soxS* induced. Of the 44 genes commonly identified in our *soxS* regulon and *E. coli* K-12 ^3^, 30 were identified to belong to the more stringent ‘oxidative *soxS*’ regulon. Our ‘oxidative *soxS* regulon’ identified many genes that were previously shown to be regulated by SoxS. A discussion of these genes is included in the supplementary file S1.

*K. pneumoniae* MGH78578 is a multi-drug resistant isolate and its antimicrobial resistance profile is well characterized ^22^. Our primary interest was in identifying how *soxS* modulates antimicrobial resistance in *K. pneumoniae* MGH78578. So we assayed whether any *K. pneumoniae* MGH78578 genes conferring antimicrobial resistance were captured in our ‘oxidative *soxS*’ regulon. We identified 11 antimicrobial resistance-encoding genes (*acrAB, acrE, tolC, marRAB, cmr* (*mdfA*), *ybhT*, KPN_RS15915, and KPN_RS15920) in the ‘oxidative *soxS’* regulon and all of them were *soxS* induced (Table S1-WS3). Interestingly we also found that another member of the XylS/AraC family, *tetD*, that was absent from the ‘oxidative regulon’ but present in the ‘*soxS*’ regulon indicating that, at least in *K. pneumoniae, tetD* is positively regulated by *soxS*. Though *tetD* was shown to modulate response against redox compounds and tetracycline ^23^, we did not find any evidence of differential regulation when *K. pneumoniae* MGH78578 was exposed to paraquat. Since many genes conferring antimicrobial resistance were modulated by *soxS*, we were interested in examining whether inactivation of *soxS* resulted in aberrations in the antimicrobial resistance pattern of *K. pneumoniae* MGH78578.

### Deletion of *soxS* in multi-drug resistant *K. pneumoniae* MGH78578 produced a significant reduction in the minimal bactericidal concentration (MBC) against antimicrobials, particularly tetracycline

The deletion of a transcriptional regulator like *soxS* could have large impacts on the cell metabolism and stress responses. To globally visualize the metabolic aberrations with respect to *soxS*, we subjected *K. pneumoniae* MGH78578 WT and its isogenic Δ*soxS* mutant to multiple growing conditions on a phenotypic microarray platform. Of the 1,484 conditions tested, altered phenotypes (WT *versus* Mutant) were observed in 517 conditions of which only 12 were significantly up-regulated wherein the mutant showed increased respiratory metabolism when compared with the WT (Supplementary file S2). Significant phenotypic alterations were found to be associated with nitrogen and some nitrogen peptides and with amino acid sources such as L-Isoleucine, L-Ornithine, and glycine. This analysis also identified 50 conditions considered to be down-regulated, in which the mutant showed reduced metabolic respiration compared with the WT. These were found to be associated with high pH (9,5) sensitivity, and having an impact on cell wall and protein synthesis due to susceptibility to antimicrobial drugs such as tetracyclines (doxycycline, demeclocycline, chlortetracycline, minocycline) aminoglycosides (amikacin), cephalosporines (cephalothin, cefuroxime, cefotaxime), β-lactams (cloxacillin, oxacillin, phenethicillin) and others such as polymyxin B and colistin (polymyxin E).

Our RNA-seq data showed that the genes encoding antimicrobial resistance such as *acrAB-tolC, marRAB*, and others, were differentially regulated in *K. pneumoniae* MGH 78578Δ*soxS* and, thus, classified as *soxS* induced. This observation, and the phenotypic microarray associated metabolic profiling, led us to hypothesize that the *soxS* mutant might have a modified antimicrobial resistance profile compared to the wild type. To test our hypothesis, we assayed the MBC of both *K. pneumoniae* MGH 78578 and *K. pneumoniae* MGH 78578Δ*soxS* against a panel of antimicrobial compounds. MBC assays were carried out on *K. pneumoniae* MGH 78578 and *K. pneumoniae* MGH 78578Δ*soxS* against colistin, gentamycin, kanamycin, cefotaxime, and tetracycline (Fig. 3). *Escherichia coli* ATCC^®^29544 was used as a control. Our results showed that there was no change in the MBC values against colistin, kanamycin, and rifampicin, even though our phenotypic microarray assay recorded a down-regulation in the metabolism of the mutant compared to the wild type. It could be that the metabolic down-regulation was not sufficient to cause an inhibitory effect. However, there was a significant reduction in the MBC values for *K. pneumoniae* MGH78578Δ*soxS* compared to *K. pneumoniae* MGH78578 when exposed to tetracycline and cefotaxime. Therefore using a combination of phenotypic microarray and RNA-seq we show that tetracycline tolerance was *soxS* dependent in MDR *K. pneumoniae* MGH78578. Oxidative stress is a common cause of cell death mediated by antimicrobial agents, irrespective of the class to which the compound belongs ^24^. So we were interested to know whether exposure to tetracycline induced any oxidative stress in *K. pneumoniae* MGH78578. For this, we checked the induction of *soxS* in tetracycline exposed *K. pneumoniae* MGH78578. Proportional induction of *soxS* expression in response to increasing tetracycline concentration confirmed that the exposure to tetracycline induced *soxS* dependent oxidative stress in *K. pneumoniae* MGH78578 (Fig. 2B).

**Fig. 3.**
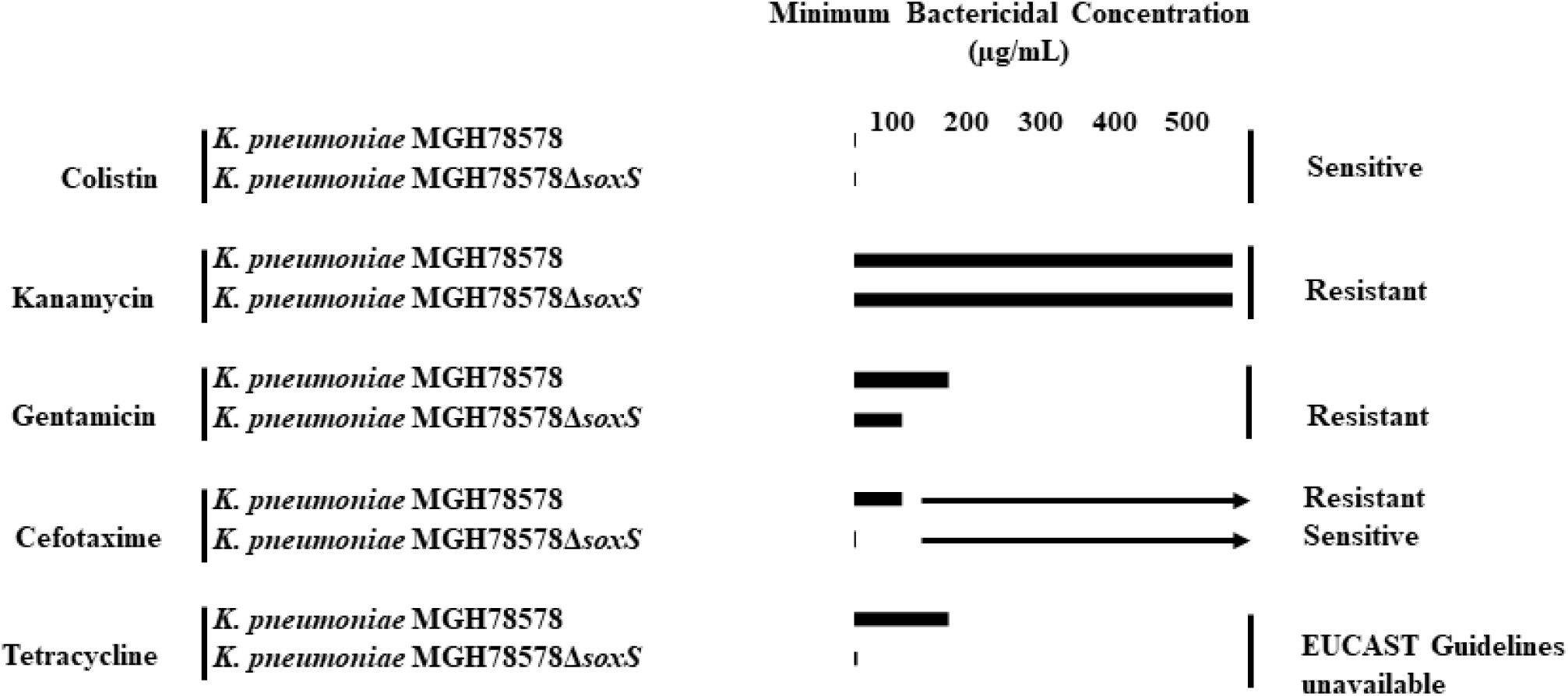
The minimum bactericidal concentrations of *K. pneumoniae* MGH78578 and *K. pneumoniae* MGH78578Δ*soxS* against different antibiotics tested.

The *soxRS* associated regulation of antibiotic resistance was described earlier in several bacteria ^25,26^. Similarly, the induction of ROS was also reported to modulate antibiotic resistance in other pathogenic bacteria. For example, *Salmonella* Typhimurium was shown to modulate its susceptibility to tetracycline when exposed to a ROS generating macrolide antibiotic, tylosin ^27^. Also, in *Acinetobacter baumannii, soxR* overexpression led to susceptibility to tetracycline ^28^. This SoxR based negative regulation of SoxS could be the reason underpinning the increased susceptibility. Even though the correlation between the expression of *soxS* and efflux pumps has been shown previously ^29^, there is no evidence pointing to the cytoplasmic accumulation of antimicrobial compounds due to an inactive *soxS* based impaired efflux activity. We, therefore, proceeded to determine whether an impaired efflux activity led to the accumulation of compounds within the cytoplasm of *K. pneumoniae* MGH78578 Δ*soxS*, leading to the bactericidal effect.

### Reduction in MBC is due to the impaired efflux pump activity in *K. pneumoniae* MGH 78578 Δ*soxS* cells

Since the *K. pneumoniae*, MGH78578Δ*soxS* was susceptible to tetracycline, we were interested in understanding the mechanism underpinning the observation. Our RNA-seq data revealed that the genes encoding the AcrAB-TolC efflux pump were highly SoxS dependent because they were >4 fold up-regulated in the PQ regulon and >8 fold down-regulated in *K. pneumoniae* MGH78578Δ*soxS*. Tetracycline is one of several structurally diverse substrates of the efflux pump AcrAB-TolC ^30^. Hence we hypothesized that the deletion of the *soxS* gene could lead to a reduction in the expression of the AcrAB-TolC efflux pump. This feature could then account for the accumulation of tetracycline in the cytoplasm to bactericidal levels.

To test our hypothesis, we assayed the efflux activity of wild type *K. pneumoniae* MGH78578 and *K. pneumoniae* MGH78578Δ*soxS* by measuring the accumulation of tetraphenylphosphonium (TPP^+^) ions using previously described protocols ^31^. We first tested whether *K. pneumoniae* MGH 78578Δ*soxS* had an intact outer membrane. In this case, both wild type *K. pneumoniae* MGH78578 and *K. pneumoniae* MGH78578 Δ*soxS* were first exposed to low concentrations of polymyxin B (PMB), an antibiotic that causes outer membrane destabilization, and then assayed the accumulation of TPP^+^. Our results showed that *K. pneumoniae* MGH78578Δ*soxS* were more susceptible to PMB and a concentration of 6 μg/ml was sufficient to induce the depolarization of the plasma membrane. In comparison, for the wild type *K. pneumoniae* MGH78578, a concentration of PMB of 9 μg/ml was required. Nonetheless, alterations in membrane voltage (maximum amount of TPP^+^) were similar for both wild type *K. pneumoniae* MGH78578 and the isogenic *K. pneumoniae* MGH78578Δ*soxS* mutant – showing that neither the outer nor the inner plasma membranes were compromised in *K. pneumoniae* MGH78578Δ*soxS* (Fig. 4A). This finding was supported by our earlier RNA-seq data which showed that membrane-associated genes that were differentially regulated during 1-(1-naphthylmethyl)- piperazine (NMP) (a chemosensitizer) treatment ^32^ were not differentially regulated in the *soxS* regulon.

**Fig. 4.**
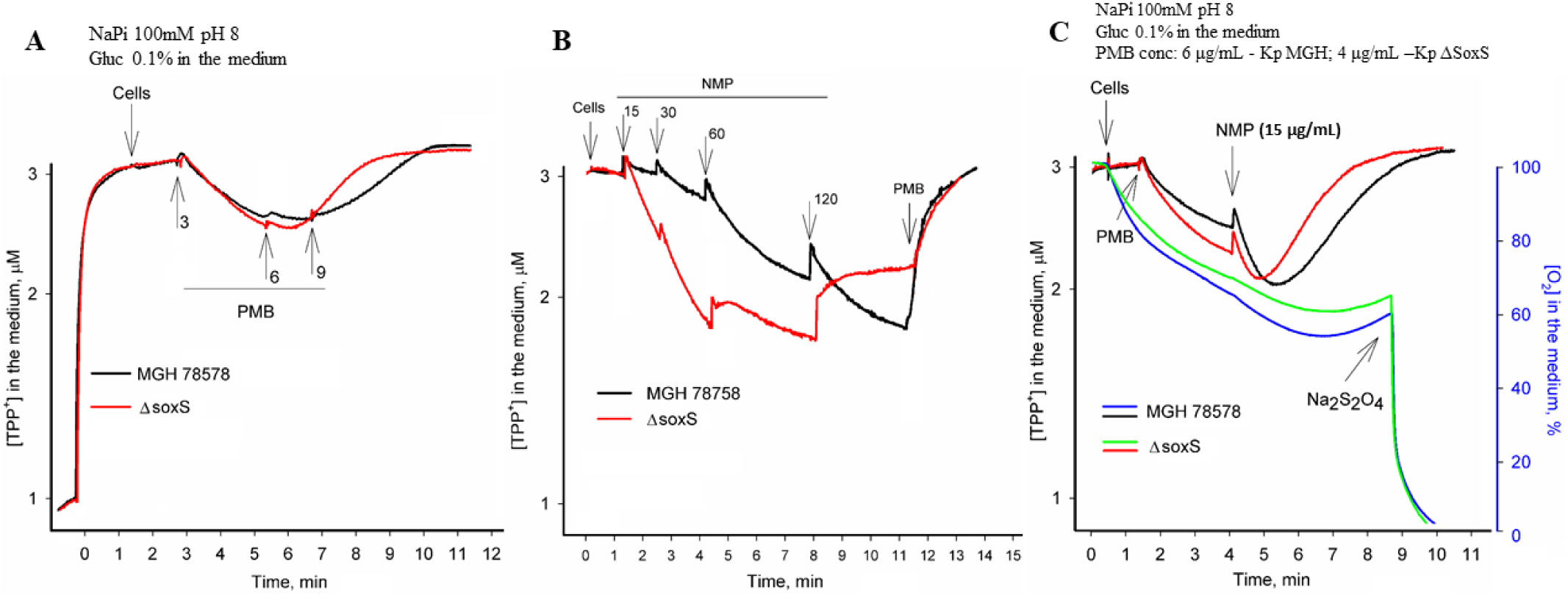
TPP^+^ accumulation in *K. pneumoniae* MGH78578 and MGH78578Δ*soxS*. The measurements were performed in 100 mM NaPi buffer containing 0.1 % glucose, pH 8.0. The concentrated cell suspensions were added to obtain OD_600 nm_ of 1. Final concentrations of NMP (μg/ml) are indicated in the figures B and C. Final concentrations of Polymixin B (PMB) are indicated in the figure (A, μg/ml), 50 μg/ml (B), or 6 and 4 μg/ml for *wt* and *ΔsoxS* cells, correspondingly (C).

Next, we investigated whether the efflux pump activity was compromised in *K. pneumoniae* MGH78578Δ*soxS* compared to that of wild type *K. pneumoniae* MGH78578. The aim was to confirm/refute our hypothesis that the impaired pump activity could result in the accumulation of tetracycline within *K. pneumoniae* MGH78578Δ*soxS*. We previously established that the treatment of *K. pneumoniae* MGH78578 with NMP destabilized the bacterial outer membrane before efflux pump inhibition and that this phenotype was concentration-dependent ^32^. Hence we used different concentrations of NMP to test the efflux pump inhibition of *K. pneumoniae* MGH78578Δ*soxS* cells compared to that of *K. pneumoniae* MGH78578. Initially, we treated wild type *K. pneumoniae* MGH78578 with NMP and assayed the cells for TPP^+^ accumulation. As expected, in wild type *K. pneumoniae* MGH78578, NMP impaired efflux pump activity and induced cytoplasmic TPP^+^ accumulation at a concenration of 30 µg/mL. However, for *K. pneumoniae* MGH78578Δ*soxS*, 15µg/mL NMP was sufficient to inhibit the efflux pump activity and cause TPP^+^ accumulation (Fig. 4B). The increased sensitivity of *K. pneumoniae* MGH78578Δ*soxS* to NMP was observed also at 120 µg/mL wherein this agent increased the accumulation of TPP^+^ in *K. pneumoniae* MGH78578 but induced a partial depolarization of the plasma membrane and leakage of the accumulated cation in *K. pneumoniae* MGH78578Δ*soxS*. These results show that the efflux pump activity was impaired in the mutant.

To further confirm the effect of the outer membrane destabilization on TPP^+^accumulation, we pre-treated both *K. pneumoniae*MGH 78578 and *K. pneumoniae* MGH78578Δ*soxS* first with PMB to permeabilize the outer membrane and then re-tested for NMP mediated TPP^+^ accumulation. These data indicated that TPP^+^ was accumulated at 15 µg/mL for *K. pneumoniae* MGH78578Δ*soxS* either indicating that both efflux pump inhibition and outer membrane destabilization could cause TPP^+^ accumulation in the bacterial cytoplasm (Fig. 4C). The respiration activity of *K. pneumoniae* MGH78578Δ*soxS* when measured was very close to that of *K. pneumoniae* MGH78578. Overall, our experiment describing the accumulation of TPP^+^ even at very low concentrations of NMP confirms that *K. pneumoniae* MGH78578Δ*soxS* exhibited an impaired efflux pump activity and as the concentration of NMP increased, membrane stability was affected resulting in TPP^+^ accumulation.

Tetracycline is a substrate for the AcrAB-TolC efflux pump. Taking together the down-regulation of *acrAB*-*tolC* and impaired efflux pump activity in *K. pneumoniae* MGH78578Δ*soxS*, we conclude that tetracycline accumulates within the cytoplasm of *K. pneumoniae* MGH78578Δ*soxS* to bactericidal concentrations making the mutant susceptible to this antibiotic.

### *K. pneumoniae* MGH78578 Δ*soxS* was avirulent in a zebrafish infection model

Since SoxS mediates oxidative stress, we sought to characterize the role of *soxS* to mitigate oxidative stress in an *in vivo* model. We used a zebrafish (*Danio rero*) embryo model to investigate the survival of *K. pneumoniae* MGH78578 Δ*soxS* in comparison with the wild type *K. pneumoniae* MGH78578. Zebrafish larvae are generally used as infection models because they are genetically tractable, optically accessible, and present a fully functional immune system with macrophages and neutrophils that mimic their mammalian counterparts ^33^. Moreover, a zebrafish larvae model was recently used to assay infection associated with *K. pneumoniae* ^34^. Bacterial strains including wild type *K. pneumoniae* MGH78578, *K. pneumoniae* MGH78578 Δ*soxS, E. coli* Xl1 blue (an avirulent control) and DPBS (un-inoculated control) were directly injected into the caudal vein of 48 hpf (hours post fertilization) zebrafish embryos and survival rates recorded by observing the presence or absence of a heartbeat post-infection. This time point was selected because the innate immune system begins to develop with primitive macrophages at 24 hpf while neutrophils develop later, at 48 hpf ^35^. Neutrophils are known to use oxidative stress to control bacterial infections in zebrafish larvae. We observed that *K. pneumoniae* MGH78578 Δ*soxS* was inefficient in killing zebrafish larvae compared to the wild type (Fig. 5). At 1 dpi (days post-infection), a survival rate of 100% was recorded in embryos injected with wild type *K. pneumoniae* MGH78578, *K. pneumoniae* MGH 78578 Δ*soxS, E. coli*, and DPBS. However, at 2 dpi, the survival rate of the embryos injected with wild type *K. pneumoniae* MGH78578 decreased to 80% while the survival rate in those embryos injected with *K. pneumoniae* MGH78578 Δ*soxS* and in the avirulent/un-inoculated controls remained unaltered. At 3 dpi, the survival rate dropped further down to 50% for the embryos injected with *K. pneumoniae* MGH78578 while being maintained at 90% for the *K. pneumoniae* MGH78578 Δ*soxS* infected embryos (Fig. 5). The survival rate remained unaltered in the case of *E. coli* and DPBS injected embryos over the time course of infection. Recent studies report that high neutrophil recruitment and zebrafish lethality is observed with *K. pneumoniae* if directly injected into the blood ^34,36^. We anticipate two possibilities for the sensitivity of *K. pneumoniae* MGH78578 Δ*soxS* in zebrafish larvae. First, in vertebrates, extracellular bactericidal action is initiated by neutrophils at a distance by activating an NADPH oxidase-dependent production of superoxide ^37^. The avirulent phenotype of *K. pneumoniae* MGH78578 Δ*soxS* could be due to the inefficiency in combating the extracellularly produced, neutrophil originated superoxide in the blood. Secondly, *K. pneumoniae* MGH78578 Δ*soxS* exhibited a down-regulated expression of *acrAB-tolC* which could result in an avirulent phenotype as seen previously in *Salmonella* Typhimurium ^38^.

**Fig. 5.**
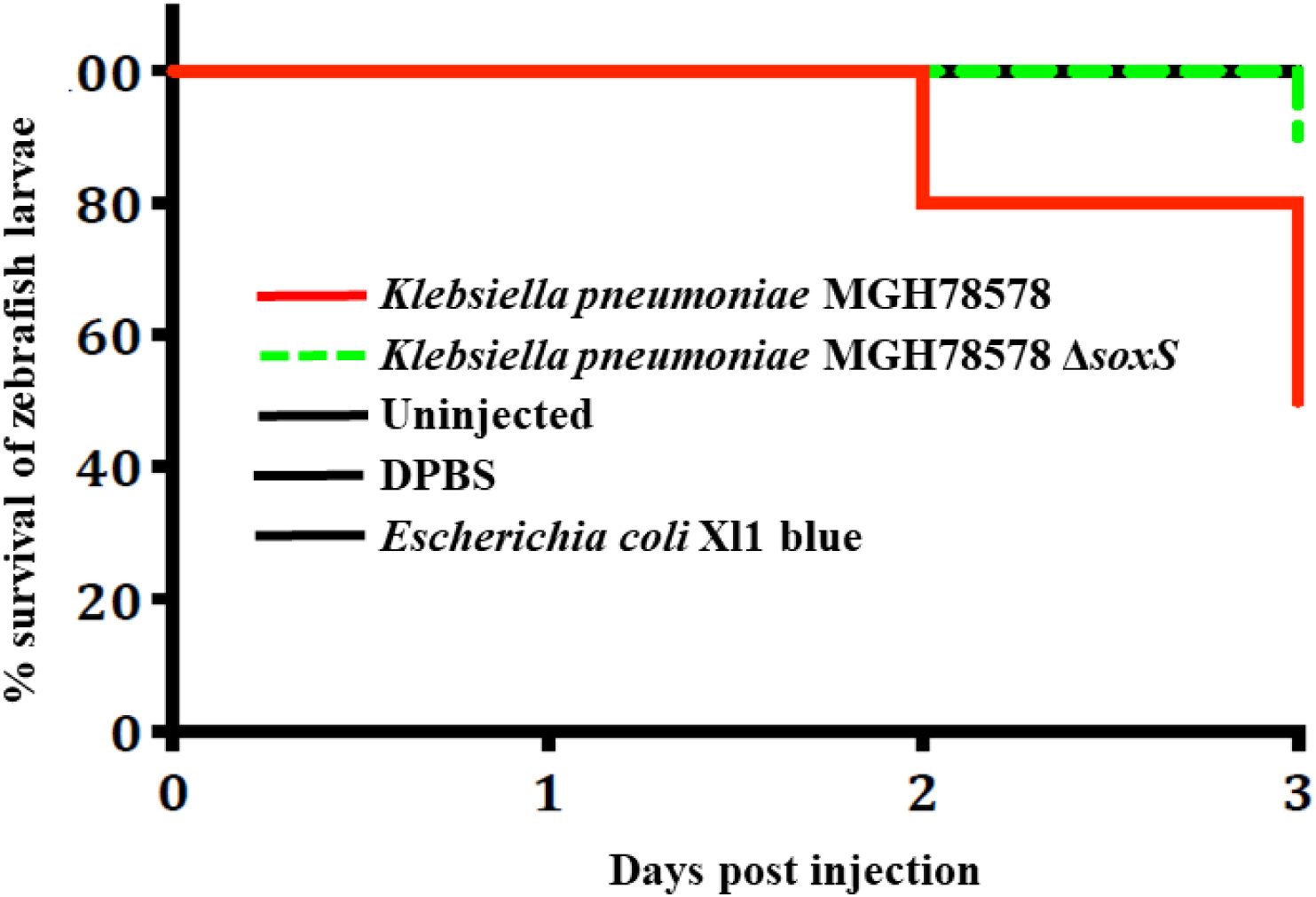
Survival of larvae in a zebrafish infection model. The survival of larvae infected with the isogenic wild type *K. pneumoniae* MGH78578 dropped as the days post-infection increased (red line). The survival of larvae infected with *K. pneumoniae* MGH78578Δ*soxS* (green line) was maintained at 100% throughout the experiment similar to the negative (avirulent *E. coli* XI1 blue, uninfected and DPBS) (black line) controls.

### By impairing *soxS*, multi-drug resistant *K. pneumoniae* MGH78578 infections can be treated by tetracycline

Currently, various strategies are being investigated to mitigate the threat of AMR in bacteria ^39^. It was thought that restricting the use of a particular antibiotic would restore susceptibility to that compound over time by eliminating the selective advantage. But it has been observed that AMR is persistent over the decades ^40^. Recent research has elaborated on the possibility wherein resistance can be reversed. One strategy here made use of defined drug-adjuvant combinations to reverse resistance so that conventional antibiotics continue to be effective ^41^. With this broad goal in mind, we endeavored to identify genetic targets that regulate resistance and develop strategies to reverse resistance by inhibiting them. Our transcriptomic and phenotypic data has shown that by inhibiting *soxS*, susceptibility to tetracycline can be restored in a multi-drug resistant *K. pneumoniae*. We were further interested to investigate whether *soxS* mediated tetracycline susceptibility can be demonstrated in an *in vivo* zebrafish model.

For this, we treated 4 hpf zebrafish embryos with increasing concentrations of tetracycline. At 48 hpf, *K. pneumoniae* MGH78578, *K. pneumoniae* MGH78578 Δ*soxS*, and DPBS were microinjected into the blood circulation. Post-injection, zebrafish larvae were collected at different time points and again bacterial counts were enumerated (Fig. 6). It was shown recently that exposure to tetracycline induced ROS production in zebrafish larvae ^42^. So, we hypothesized that *K. pneumoniae* MGH78578 Δ*soxS* will be impaired in its ability to survive in tetracycline treated zebrafish larvae due to increased sensitivity to either ROS production in tetracycline treated larvae or tetracycline alone. It is also possible that healthy zebrafish larvae could clear *K. pneumoniae* MGH78578 Δ*soxS* from the system due to normal exposure to peroxides synthesized from neutrophils. Confirming our hypothesis, *K. pneumoniae* MGH78578 Δ*soxS* was completely cleared from the tetracycline treated zebrafish larvae in 24 hours. However, *K. pneumoniae* MGH78578 Δ*soxS* was cleared even in tetracycline untreated zebrafish larvae confirming that the selective advantage was lost in the bacterial mutant making it susceptible to the immune system of zebrafish larvae (Fig. 6). It should also be noted that the clearance was much more marked in tetracycline treated larvae suggesting that *K. pneumoniae* MGH78578 Δ*soxS* was cleared from the system due to a cumulative effect of both immune system and tetracycline induced ROS production. Overall, we show that *soxS* can be used as a genetic target to treat multi-drug resistant *K. pneumoniae* infections.

**Fig. 6.**
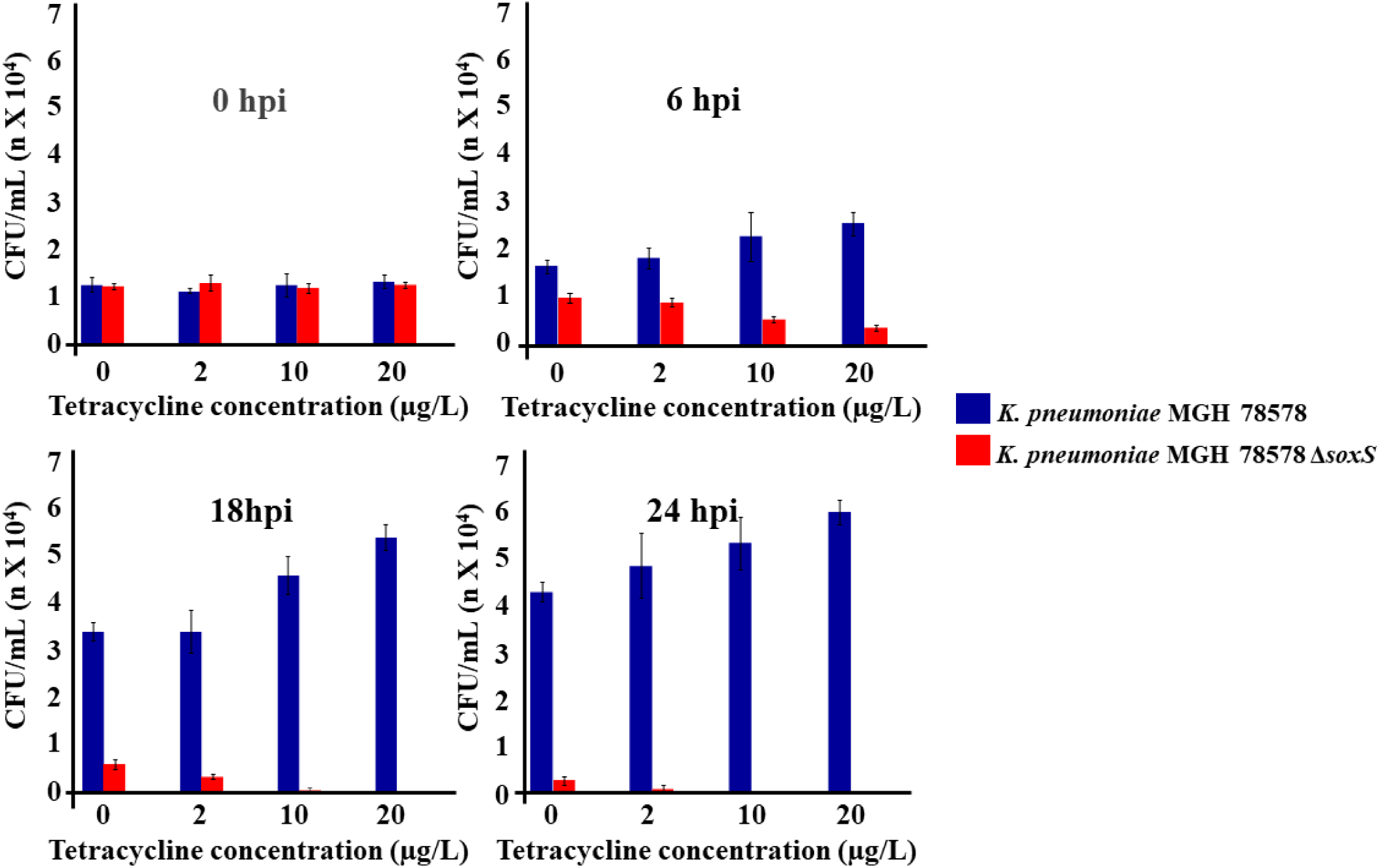
Survival of bacteria in a zebrafish larvae infection model where the larvae are treated with tetracycline. Tetracycline induces ROS generation in the zebrafish larvae. The bar charts show the survival of the different bacterial cultures in the larval blood at 0, 6, 18, and 24 hours post-infection, and at different concentrations of tetracycline.

### Conclusion

Apart from elucidating the PQ oxidative stress regulon and the oxidative SoxS regulon, we propose that a combination of tetracycline and a SoxS inhibitor can be used to treat infections associated with MDR *K pneumoniae*. Tetracycline will induce oxidative stress in the host and the SoxS inhibitor can impair the ability of the pathogen to survive oxidative stress. The advantage of this approach is that it is independent of any particular bacterial resistance mechanism, can be used against strains with any resistance profile. Certainly, the next step in this approach is to construct/identify a molecule that will specifically and post-transcriptionally inhibit the synthesis of *soxS* mRNA or post-translationally inhibit the SoxS protein. It will be worthwhile to test if our observations will also apply to other MDR pathogens like *Salmonella* Typhimurium, *E. coli* among others. Nevertheless, our results address the immediate concern of antimicrobial resistance in pathogens of importance to human health whilst providing a ‘proof of concept’ for an approach that requires further experimental investigation to achieve the therapeutic objective.

## Materials and methods

### Bacterial strain

MDR *Klebsiella pneumoniae* MGH78578 (ATCC^®^700721) was isolated from a sputum sample in 1994 and was purchased from the American Type Culture Collection. This strain was selected mainly because it is a multi-drug resistant type strain ^43^ and its drug resistance profile was recently published ^22^. Moreover, the efflux pumps present in this strain are well characterized ^44,45^. Further, the whole genome sequence of this strain is available in NCBI (Reference Sequence NC_009648.1) which was convenient for mapping RNA-seq data. This bacterium was grown in Müeller-Hinton (MHB) broth and Müeller-Hinton agar (MHA) (Sigma, Dublin, Ireland).

### Phenotypic assay (OmniLog)

The comparison of *K. pneumoniae* MGH78578 WT with the Δ*soxS* mutant was evaluated using the OmniLog (Biolog, Inc., Hayward, CA) phenotypic microarray. Microplates PM1 through PM20 with the exception of PM5 were used. These plates contain several carbon, nitrogen, sulphur and phosphorous-substrates, ions, osmolytes and chemicals at different concentrations and pH ^46^. *Klebsiella pneumoniae* MGH78578 WT and Δ*soxS* were grown at 37°C on LB agar plates, and several colonies were picked with a sterile cotton swab and suspended in 15 mL IF-0 until a cell density of 42% transmittance (T) was reached (measured using a Biolog turbidimeter). Each 15 mL suspension was then added to 75 mL of physiological solution IF-0 containing dye A used to inoculate PM plates 1-2. Plates PM 3-8 were inoculated with IF-0 solution containing sodium pyruvate as carbon source. PM 9-20 were inoculated with the physiological solution IF-10. One hundred microliters of each mixture were inoculated into each well of the microplates. All PM microplates were incubated at 37°C in an OmniLog reader and monitored for 72 h. Data was analysed using DuctApe software v 0.17.4 ^47^. Each strain was analysed in duplicate. Results are present in supplementary file S2.

### Isolation of RNA from oxidatively stressed bacterial cells

Before RNA isolation, wild type *Klebsiella pneumoniae* MGH78578 and *K. pneumoniae* MGH78578 Δ*soxS* were grown until mid-exponential state (MEP) following an earlier standardized protocol ^22^. MEP grown bacterial cells were treated with paraquat (7.81 µM) for 30 minutes to generate oxidative stress conditions. RNA was then extracted from both oxidative stressed and MEP grown (control) cells using QIAGEN RNeasy Mini Kit following manufacturers guidelines. Contaminating DNA was removed from the RNA sample using the Turbo DNase I kit (Thermo Fischer Scientific). RNA was then quantified using both Qubit RNA Broad Range Assay and the Nanodrop.

### Sequencing RNA isolated from oxidatively stressed bacterial cells

The library preparation and subsequent sequencing were carried out commercially at the Center For Genomic Research, University of Liverpool. Ribo-Zero rRNA removal Kit for Bacteria (Illumina, San Diego, CA) was used to carry out the depletion of ribosomal DNA according to the manufacturer’s instructions. Libraries were created using NEBNext® Ultra(tm) Directional RNA Library Prep Kit (New England BioLabs, Frankfurt, Germany). Pooled libraries were loaded on the cBot (Illumina, San Diego, CA) and cluster generation was performed according to the manufacturer’s instructions. Single-end sequencing using 125 bp read length was performed on an Illumina HiSeq 2500 platform (Illumina HiSeq Control Software 2.2.38) using an Illumina HiSeq Flow Cell v4 and TruSeq SBS Kit v4 (Illumina). Raw sequencing read data was processed using RTA version 1.18.61 and CASAVA 1.8.4 to generate FASTQ-files. Genomic cDNA libraries were prepared using the TruSeq Stranded Total RNA Library Prep Kit (Illumina, San Diego, CA) with Ribo-Zero to deplete ribosomal RNA (rRNA). An average of 1.48 Gbp of raw sequence data was obtained per sample, in 125 bp single-end reads.

### Mapping of sequenced reads

The sequence quality of the RNA-seq reads was analyzed using FastQC [https://www.bioinformatics.babraham.ac.uk/projects/fastqc/]. Sequence reads were aligned and mapped against the reference genome of *K. pneumoniae* MGH78578 (Reference Sequence NC_009648.1) using Segemehl with default mapping parameters ^22,48^ and uniquely mapped reads were used and considered for the differential gene expression computational analysis. Read counts (number of reads that aligned to a specific gene) for each gene were quantified using custom Perl scripts.

### Computational analysis of RNA-seq data

Computational analysis of RNA-seq data was performed using R (version 3.5.2, https://www.r-project.org/). To calculate the expression level of genes, the raw read counts were normalized using the VOOM function ^21^ in the limma package ^49^. More specifically, counts were converted to log_2_ counts per million (log_2_ CPM), quantile normalized, and precision weighted using the VOOM function. A linear model was then fitted to each gene, empirical Bayes moderated t-statistics and its corresponding p-values were used to assess differences in expression ^21,50^. To account for multiple comparisons, Benjamini-Hochberg corrected p-values were computed. As reads for duplicated coding genes (paralogs) or duplicated small RNAs cannot be mapped unequivocally, these genes appear in the analysis as unmapped. The sequence reads can be visualized in the Integrated Genome Browser (version 9.0.0) _51_. The read depth was adjusted with the cDNA library with the lowest number of reads ^52^. RNA sequencing data were analyzed using the fold change parameters as follows: highly up-regulated (>4), up-regulated (2 to 4 fold), no change in expression (0.5 to 2 fold), down-regulated (0.25 to 0.5 fold) and highly down-regulated (less than 0.25 fold).

### Construction of the *K. pneumoniae* MGH78578 Δ*soxS* mutant

A modified λ-Red system was used to construct an in-frame deletion in multi-drug resistant *K. pneumoniae* MGH78578 ^53^. Here three plasmids are employed. The first, plasmid pIJ773 which serves as a template to amplify the apramycin resistance gene, *aac(3)IV*, and flanking FRT sites. The second plasmid, pACBSR-Hyg contains the λ-Red system comprising *beta, gam*, and *exo* genes which are under the control of an arabinose-inducible promoter and facilitate homologous recombination between the knockout cassette and the target locus in the chromosome. The third plasmid, pFLP-Hyg, contains the FLP recombinase, which was used to excise the apramycin selection marker from the chromosome via the FRT sites. The antibiotic apramycin was used to select plasmid pIJ773, while hygromycin was used to select both plasmids pACBSR-Hyg and pFLP-Hyg. *K. pneumoniae* MGH78578 was susceptible to both apramycin and hygromycin.

Plasmid pACBSR-Hyg was first introduced into wild-type *K. pneumoniae* MGH78578 by electroporation. An overnight culture of the bacteria was reinoculated into 100 mL Luria Bertani (LB) broth at 220 rpm aeration and 30°C temperature until an OD_600 nm_ of 0.6 to 0.8 was reached. The cells were washed twice with 50 ml ice-cold 10% (v/v) glycerol and resuspended in the residual glycerol solution after the final wash. A 50 μl aliquot of the dense suspension was then mixed with 200 to 400 ng of plasmid DNA and electroporated at 2,500 mV. Bacterial cells were revived in SOC medium which was added immediately after electroporation and *K. pneumoniae* MGH78578 [pACBSR-Hyg] were selected by plating on low salt LB plates containing hygromycin. The knockout cassette consisting of three distinct regions: the *aac(3)IV*, FRT sites that flanked *aac(3)IV*, and 60 bp regions homologous to the *soxS* gene were amplified using PCR from plasmid pIJ773 template as the template. Competent *K. pneumoniae* MGH78578 [pACBSR-Hyg] were prepared by growing the cells grown in low salt LB with 1 M L-arabinose and 100 µg/mL hygromycin – and washing in ice-cold 10% (v/v) glycerol as described above. The PCR amplified knock out cassette was electroporated into competent *K. pneumoniae* MGH78578 [pACBSR-Hyg] and cells with successful recombination events were selected by plating in LB with apramycin and incubating overnight at 37°C. *K. pneumoniae* MGH78578 *soxS*::FRT-*aac(3)IV*-FRT cells were identified and confirmed using PCR primers targeting regions that flanked *soxS* region. Competent *K. pneumoniae* MGH78578 *soxS*::FRT-*aac(3)IV*-FRT cells were prepared and electroporated with plasmid pFLP-Hyg to excise the inserted knock out cassette and the *K. pneumoniae* MGH78578 Δ*soxS* cells were confirmed using PCR and sequencing. The sequences of all primers used in the experiment is provided in Table S1 WS-1.

### Isolation of RNA for Quantitative Reverse Transcriptase Polymerase Chain Reaction (qRT-PCR)

Wild type *K. pneumoniae* MGH78578 was grown to Mid Exponential Phase (MEP) at 37°C in Müeller-Hinton Broth, cells were then treated with paraquat at different concentrations (0, 3.905, 7.81, 200 and 500 µM) or similarly with tetracycline (0, 0.5, 10, 100 500 μg/mL) for 30 minutes and RNA was then extracted. All assays were run in triplicate. In all conditions, RNA was extracted using QIAGEN RNeasy Mini Kit following manufacturers guidelines. Any contaminating DNA was removed from the RNA sample using the Turbo DNase I kit (Thermo Fischer Scientific). Purified RNA was subsequently quantified using both Qubit RNA Broad Range Assay and Nanodrop.

### Two steps Quantitative Reverse Transcriptase Polymerase Chain Reaction (qRT-PCR)

The reverse transcriptase reaction was carried out on RNA purified from *K. pneumoniae* MGH78578 under the conditions mentioned earlier, using the high capacity RNA to cDNA kit (Sigma) following manufacturer guidelines. A negative control devoid of RT enzyme was also carried included. qPCR was then performed following the prime-time gene expression master mix protocol (IDT, Leuven). The expression of the housekeeping gene *rpoB* was used to characterize the relative expression of the gene of interest, *soxS*. The sequences of all primers used in the experiment is provided in Table S1 WS-1.

### Determination of Minimum Bactericidal Concentration (MBC)

Previously, MIC values for Paraquat (PQ), colistin (COL), tetracycline (TET), gentamicin (CN), kanamycin (KM) and cefotaxime (CTX) were determined in triplicate using a 96-well microtiter plate two-fold broth microdilution method ^22^. The range of concentrations employed was 1-512 µg/mL for all antibiotics excluding colistin for which a concentration range of 0.03125-16 µg/mL and paraquat 0.97-500µM was utilized. Overnight LB cultures of *K. pneumoniae* MGH78578 and *K. pneumoniae* MGH78578 Δ*soxS* were diluted in sterile phosphate buffered saline to 10^5^ CFU/mL concentration. A 96-well plate was used to prepare two-fold serial dilutions of each antibiotic for MHB and MBC determination of *K. pneumoniae* MGH78578 and *K. pneumoniae* MGH78578 *ΔsoxS* against each compound. A volume of 5 µL of the 10^5^ CFU/mL bacterial culture was then transferred to separate wells containing varying concentrations of the compounds to be tested. These plates were then incubated at 37°C for 16-18 hours according to the Clinical Laboratory Standards Institute (CLSI) guidelines. Triplicate MBC values for each antibiotic tested were determined using MHB broth in a 96-well microtiter plate. A steel inoculator was employed to transfer inoculum from the above 96-well plate as described above to a fresh 96-well plate containing MHB without any of the antibiotics to be tested. These plates were then incubated at 37°C for 16-18 hours, following which the MBC values were recorded.

### Electrochemical measurement experiments

The efflux activity of *K. pneumoniae* MGH78578 and *K. pneumoniae* MGH78578 Δ*soxS* cells was assayed measuring accumulation of tetraphenylphosphonium (TPP^+^) ions. Over-night cultures of *K. pneumoniae* were grown in Luria-Bertani broth, containing 0.5 % NaCl, diluted 3:100 in fresh medium, and the incubation was continued until the OD_600 nm_ reached 1.0. The cells were collected by centrifugation at 4 °C for 10 min at 3000 g. The pelleted cells were re-suspended in 100 mM sodium phosphate buffer, pH 8, to obtain 1,4 × 1011 cfu/ml. Concentrated cell suspensions were kept on ice until used, but not longer than 3 h.

Changes of TPP^+^ concentration in the suspensions of thermostated and magnetically stirred cells were monitored using TPP+-selective electrodes as previously described ^31 54^. Experiments were performed at 37 °C in 100 mM sodium phosphate buffer, pH 8, containing 0.1 % glucose. OD_612 nm_ of the cell suspension during measurements was 1.

### Zebrafish line maintenance, infection and microinjection experiments

Zebrafish (*Danio rerio*) used in this study were *wik* lines. Adult fish were kept at a 14/10 h light/dark cycle at a pH of 7.5 and 27°C. Eggs were obtained from natural spawning adult fish which were set up pairwise in individual breeding tanks. Embryos were raised in Petri dishes containing E3 medium (5 mM NaCl, 0.17 mM KCl, 0.33 mM CaCl_2_, 0.33 mM MgSO_4_) supplemented with 0.3 µg/ml of methylene blue at 28°C. From 24 hpf, 0.003% 1-phenyl-2-thiourea (PTU) was added to prevent melanin synthesis. The staging of embryos was performed as explained earlier ^55^.

Microinjections were performed using borosilicate glass microcapillary injection needles (Science Products, 1210332, 1 mm O.D. x 0.78 mm I.D.) and a PV830 Pneumatic PicoPump (World Precision Instruments). The 48 hpf embryos were manually dechorionated and anesthetized with 200 mg/l buffered tricaine (Sigma, MS-222) before injection. Subsequently, embryos were aligned on an agar plate and injected with 12000 CFU in 1-4 ml volume of a bacterial suspension in DPBS directly into the blood circulation (caudal vein, n=10 x3). Before injection, the volume of the injection suspension was adjusted by injecting a droplet into mineral oil and measuring its approx. diameter over a micrometer scale bar. The number of CFU injected was determined by injection of bacterial suspension into a DPBS droplet on the agar plate. Following injections, injected embryos were allowed to recover in a petri dish with fresh E3 medium for 15 min. To follow infection kinetics and for survival assays, embryos were transferred into 24-well plates (one embryo per well) in 1 ml E3 medium per well, incubated at 28°C and observed for signs of disease and survival under a stereomicroscope twice a day. For survival assays after infection, the number of dead larvae was determined visually based on the absence of a heartbeat. Kaplan Meier survival analysis and statistics for experiments with Zebrafish was done with GraphPad Prism 7 (GraphPad Software, United States). Experiments were performed in triplicate.

## Data availability

All the RNA sequence data generated in the study are deposited in the National Center for Biotechnological Information – Gene Expression Omnibus and are available under the accession number GSE146844. Post analysis, the differential expression of all the genes are given in supplementary Table S1, with three worksheets WS-1, WS-2 and WS-3.

**Table S1-WS1**: RNA-seq mapping details. Details of the RNA-seq reads mapped against different regions of the *K. pneumoniae* MGH78578 genome is given here. Is also given here a list of all primers used in the experiments in this paper.

**Table S1-WS2:** The gene expression pattern of *Klebsiella pneumoniae* MGH75878 (MGH_PQ_ *versus* MGH_WT_) and *K. pneumoniae* MGH75878 Δ*soxS* (MGH *Δsox*S_PQ_ *versus* MGH_PQ_) during oxidative stress. Column A shows a new gene ID and column C shows Old name while column B represents the gene name. Column X/Y/Z/AA/AB/AC/AD shows raw reads generated from RNA-seq conducted on different libraries while column AF/AG/AH/AI/AJ/AK represents the normalized reads. The fold changes were calculated from the normalized reads obtained from different libraries MGH_PQ_ *versus* MGH_WT_ (column K) MGH Δs*oxS*_*PQ*_ *versus* MGH_PQ_ (column U). The genes with statistically significant data obtained from two biological replicates are indicated by √ mark in column J and T based on p-value indicated in column M and W. Based on the differential expression indicated in column K and U (for statistically significant genes), the differential expression of each gene is indicated by a color code in column E/F/G/H and O/P/Q/R. Genes highlighted in red indicate antimicrobial resistance genes while those highlighted in blue indicate genes associated with virulence.

**Table S1-WS3:** The ‘oxidative *soxS* regulon’ of *Klebsiella pneumoniae* MGH78578 - a stringent set of genes that are regulated by SoxS. The gene list was obtained from statistically significant genes from oxidative regulon and *soxS* regulon with a distinct expression pattern. Genes those were up-regulated in the oxidative regulon + down-regulated in *soxS* regulon and down-regulated in oxidative regulon + up-regulated in *soxS* regulon.

## Code availability

All codes used are published programs, with citations for each provided in the references.

## Acknowledgment

We gratefully acknowledge Enterprise Ireland (IP 2015 0380) for funding JA and SS. JA would also like to acknowledge the financial support through the research grant 11/F/051 provided by the Department ofAgriculture, Food and the Marine (DAFM), Ireland. The funders had no role in study design, data collection, and analysis, decision to publish, or preparation of the manuscript.

## Author Contributions

JA, SF, and SS conceived and supervised the experiments. JA, KD, and YC performed the growth, PQ induction, RNA-seq, and qRT – PCR experiments. SKS and SN carried out the bioinformatics analysis of the RNA-seq datasets. SaS and RD carried out the TPP^+^ assays. AE and AL carried out the zebrafish embryo assays. JA, SD, SF, and SS wrote this publication into which all other authors contributed. All authors read and approved the manuscript.

## Competing Interests

The authors declare no competing interests.

## Notes

### Competing Interest Statement

The authors have declared no competing interest.

