## Supplementary files S1 for "Analysis of the oxidative stress regulon identifies *soxS* as a genetic target for resistance reversal in multi-drug resistant *Klebsiella pneumoniae*"

**Supplementary file S1**

A report from 1990 used *lac* fusions to identify five PQ inducible promoters (including *sodA*, *nfo*, *zwf*, *fumC*, and *fpr*) in *E. coli* (Greenberg et al. 1990), all of which played protective roles during PQ mediated oxidative stress. All the above-mentioned genes were identified in our 'oxidative *soxS*' regulon and were 'SoxS induced' (Table S1-WS3). The gene *sodA* encodes a manganese superoxide dismutase which will convert two molecules of superoxide to oxygen and hydrogen peroxide (Broxton and Culotta 2016). The latter is then removed by catalase and peroxide enzymes. The gene encoding *sodA* was up-regulated and down-regulated ~15 times respectively in $MGH_{par}$ and MGHΔ*soxS*$_{par}$ library – demonstrating the protective nature of *sodA* during oxidative stress. Exposure to oxidative stress can cause lesions in DNA which are then repaired by the base excision repair (BER) pathway. In *E. coli* endonuclease IV encoded by *nfo* is involved in this BER pathway enabling repair of DNA lesions associated with oxidative stress (Souza et al. 2006). The gene *nfo* was up-regulated ~14 times in the $MGH_{par}$ and was down-regulated ~ 20 times in MGH Δ*soxS*$_{par}$ library – confirming the importance of *nfo* in the defence against oxidative stress. We also noted an ~ 7f- up-regulation and ~ 4 f- down-regulation of the gene *zwf* encoding a glucose 6-phosphate dehydrogenase, respectively in the $MGH_{par}$ and MGHΔ*soxS*$_{par}$ library (Table S1-WS2). Up-regulation of *zwf* results in increased synthesis of reduced nicotinamide adenine dinucleotide phosphate (NADPH) which is critical in maintaining a balanced redox status and improved survival of *K. pneumoniae* during oxidative stress, as reported earlier in *E. coli* (Ying 2008; Sandoval et al. 2011). The gene *fumC* encodes a fumarase C which functions in combating oxidative stress and which was first noted in *E. coli* (Liochev and Fridovich 1992). As exemplified in *E. coli*, fumarases (three in total - encoded by *fumA, fumB* and *fumC*) play an important role in the citric acid cycle. While *fumA* and *fumB* are important in anaerobic growth, *fumC* had the highest activity during growth in aerobic conditions (Tseng et al. 2001). In *E. coli* the activity of *fumC* is to maintain the activity of fumarases during oxidative stress since highly oxidative conditions inactivate *fumA*. The up-regulation of *fumC* to ~37f- in *K. pneumoniae* MGH78578 and down-regulation to ~20f- in *K. pneumoniae* MGH78578 Δ*soxS* could be to maintain the activity of citric acid cycle even under extreme oxidative stress since fumarase C is resistant to oxidative conditions (Tseng et al. 2001). However, no homologues of *fumA* nor *fumB* were identified in the genome of *K. pneumoniae* MGH78578.

Another member of the *K. pneumoniae* MGH78578 'oxidative *soxS* regulon', *ydbK*, was up-regulated ~ 48f- in the $MGH_{par}$ and down-regulated ~50f- in MGHΔ*soxS*$_{par}$ (Table S1-WS3). Similarly, *fpr*, encoding a ferredoxin (flavodoxin)-NADPH reductase was also up-regulated ~35f- and down-regulated ~20f-. YdbK is a pyruvate:flavodoxin oxidoreductase and is susceptible to oxygen. In *E. coli*, the gene *ydbK* was overexpressed during exposure to superoxide generators like methyl viologen in a SoxS dependent manner (Nakayama et al. 2013). An *E. coli* double mutant of *ydbK* and *fpr* was significantly sensitive to oxidative stress compared to individual mutants alone (Nakayama et al. 2013). Our RNA-seq data shows that in *K. pneumoniae*, *ydbK* and *fpr* play antioxidant roles in independent but related pathways, as shown in *E. coli* earlier. In *E. coli* too, the genes *nfsA* and *yjeC* form an operon that can be induced by PQ and regulated by *soxS* (Paterson et al. 2002). NfsA is an oxygen insensitive nitroreductase that mediates single electron transfers from NADPH to molecular oxygen. The process generates nitro-anion free radicals which are re-oxidised to generate superoxides and is one of the *soxS* mediated mechanisms behind superoxide production during bacterial exposure to paraquat (Peterson et al. 1979). Similar to *E. coli*, our *K. pneumoniae* MGH78578 'oxidative *soxS*' regulon identified *nfsA* as up-regulated during exposure to paraquat (>37 fold) and down-regulated ~33 fold in *K. pneumoniae* MGH78578 Δ*soxS* (Table S1-WS3). As early as in 1990, the gene *ompF* was found to the down-regulated during exposure of *E coli* to PQ mediated oxidative stress (Greenberg et al. 1990). In *K. pneumoniae* MGH78578 too, *ompF* showed a 2f-down-regulation in $MGH_{par}$ and a 2f- up-regulation in $MGH\Delta soxS_{par}$ cells.

### Supplementary figures

**Figure S1.** Distribution of RNA-seq reads in each individual dataset.

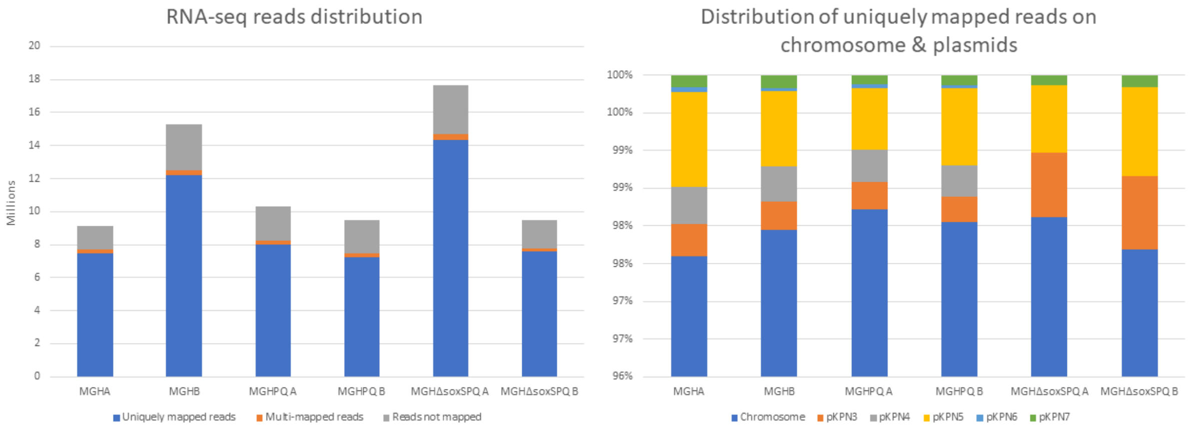
